# Microvasculopathy And Soft Tissue Calcification In Mice Are Governed by Fetuin-A, Pyrophosphate And Magnesium

**DOI:** 10.1101/577239

**Authors:** Anne Babler, Carlo Schmitz, Andrea Büscher, Marietta Herrmann, Felix Gremse, Theo Gorgels, Jürgen Floege, Willi Jahnen-Dechent

## Abstract

**Objective:** Calcifications can disrupt organ function in the cardiovascular system and the kidney, and are particularly common in patients with chronic kidney disease (CKD). Fetuin-A deficient mice maintained against the genetic background DBA/2 exhibit particularly severe soft tissue calcifications, while fetuin-A deficient C57BL/6 mice remain healthy. We employed molecular genetic analysis to identify risk factors of calcification in fetuin-A deficient mice. We sought to identify pharmaceutical therapeutic target that could be influenced by dietary of parenteral supplementation.

**Approach and Results:** We studied the progeny of an intercross of fetuin-A deficient DBA/2 and C57BL/6 mice to identify candidate risk genes involved in calcification. We determined that a hypomorphic mutation of the *Abcc6* gene, a liver ATP transporter supplying systemic pyrophosphate, and failure to regulate the TRPM6 magnesium transporter in kidney were associated with severity of calcification. Calcification prone fetuin-A deficient mice were alternatively treated with dietary phosphate restriction, magnesium supplementation, or by parenteral administration of fetuin-A or pyrophosphate. All treatments markedly reduced soft tissue calcification, demonstrated by computed tomography, histology and tissue calcium measurement.

**Conclusions:** We show that pathological ectopic calcification in fetuin-A deficient DBA/2 mice is caused by a compound deficiency of three major extracellular and systemic inhibitors of calcification, namely fetuin-A, pyrophosphate, and magnesium. All three of these are individually known to contribute to stabilize protein-mineral complexes and thus inhibit mineral precipitation from extracellular fluid. We show for the first time a compound triple deficiency that can be treated by simple dietary or parenteral supplementation. This is of special importance in patients with advanced CKD, who commonly exhibit reduced serum fetuin-A, pyrophosphate and magnesium levels.

**Subject Codes:** Animal Models of Human Disease, Fibrosis, Inflammation, Proteomics, Peripheral Vascular Disease

## Introduction

Pathological calcifications, the gradual deposition of calcium phosphate salts [2] in tissue not physiologically meant to mineralize are frequent and are mostly considered benign. However, in particular in the context of chronic kidney disease (CKD), vascular calcifications have increasingly been recognized as a major contributor to cardiovascular morbidity and mortality independent of traditional risk factors [3].

Cell autonomous signaling and the recruitment of mesenchymal cells promote the progression of late stage calcifications to ossifications [4] mimicking bone formation in soft tissues [5]. In most cases, however, calcifications start in the extracellular matrix by nucleation of calcium phosphate crystals in the absence of mineralization regulators [6], before any osteogenic reprograming of resident or invading mesenchymal cells occurs [7, 8].

Phosphate retention in CKD is as a major driver of vascular calcification, endothelial damage, and the cardiovascular morbidity and mortality associated with CKD [6]. A disturbed phosphate homeostasis is closely associated with calcifications and accelerated ageing [9]. Consequently, dietary and blood phosphate reduction are prime targets of renal replacement therapy.

Another risk factor for the development of extraosseous calcifications especially in CKD patients is a reduced level of the serum protein fetuin-A [10]. Lack of fetuin-A allows spontaneous mineral nucleation and growth, and hydroxyapatite crystals are deposited causing cardiovascular calcifications [11] and possibly also calciphylaxis [12].

Additionally, ectopic calcification is prevented by low molecular compounds like pyrophosphate [13], and magnesium [14], which inhibit formation of hydroxyapatite through inhibition of crystal nucleation. Both have been reported to be reduced in sera of patients with advanced CKD [15, 16].

Several years ago we generated mice with a severe spontaneous soft tissue calcification phenotype [17] that worsens progressively throughout life. The mice are deficient in the liver-derived plasma protein fetuin-A, and are maintained against the genetic background DBA/2, which even in the presence of fetuin-A is prone to dystrophic cardiac calcifications [18, 19]. Fetuin-A deficiency in DBA/2 mice greatly worsens the calcification propensity, and is associated with decreased breeding performance and increased mortality. In addition, severe renal calcinosis causes CKD and secondary hyperparathyroidism [17]. Ultimately in these mice, myocardial calcification is associated with fibrosis, diastolic dysfunction and increased mortality [20, 21]. In contrast fetuin-A deficient mice maintained against the genetic background C57BL/6 do not calcify spontaneously. However, in a model of CKD induced via nephrectomy and high dietary phosphate diet, these mice developed cardiovascular calcifications [22] mimicking the situation in CKD patients.

Here we studied the calcification in progeny of an intercross of fetuin-A deficient DBA/2 and C57BL/6 mice to identify molecular determinants of their differential calcification that might serve as therapeutic targets.

## Results

### Severe ectopic calcification in fetuin-A deficient mice

We performed computed tomography (CT) in wildtype and fetuin-A deficient DBA/2 and C57BL/6 mice. The respective phenotypes were confirmed by Western blot analysis (suppl. Figure 1). Figures 1a-d show typical CTs of mice at age 3 months. Figures 1a,b illustrate that DBA/2 wt mice had rare, if any lesions in their interscapular brown adipose tissue, and C57BL/6 wt mice had no calcified lesions. In contrast, DBA/2 fetuin-A deficient mice had numerous and widespread calcifications of their brown adipose tissue, the skin rubbing against the humerus, the heart and the kidneys (Fig. 1c). Unlike the strongly calcifying DBA/2 mice, age-matched C57BL/6 mice had few if any calcified lesions in their interscapular brown adipose tissue confirming their relative calcification resistance (Fig. 1d).

**Figure 1.**
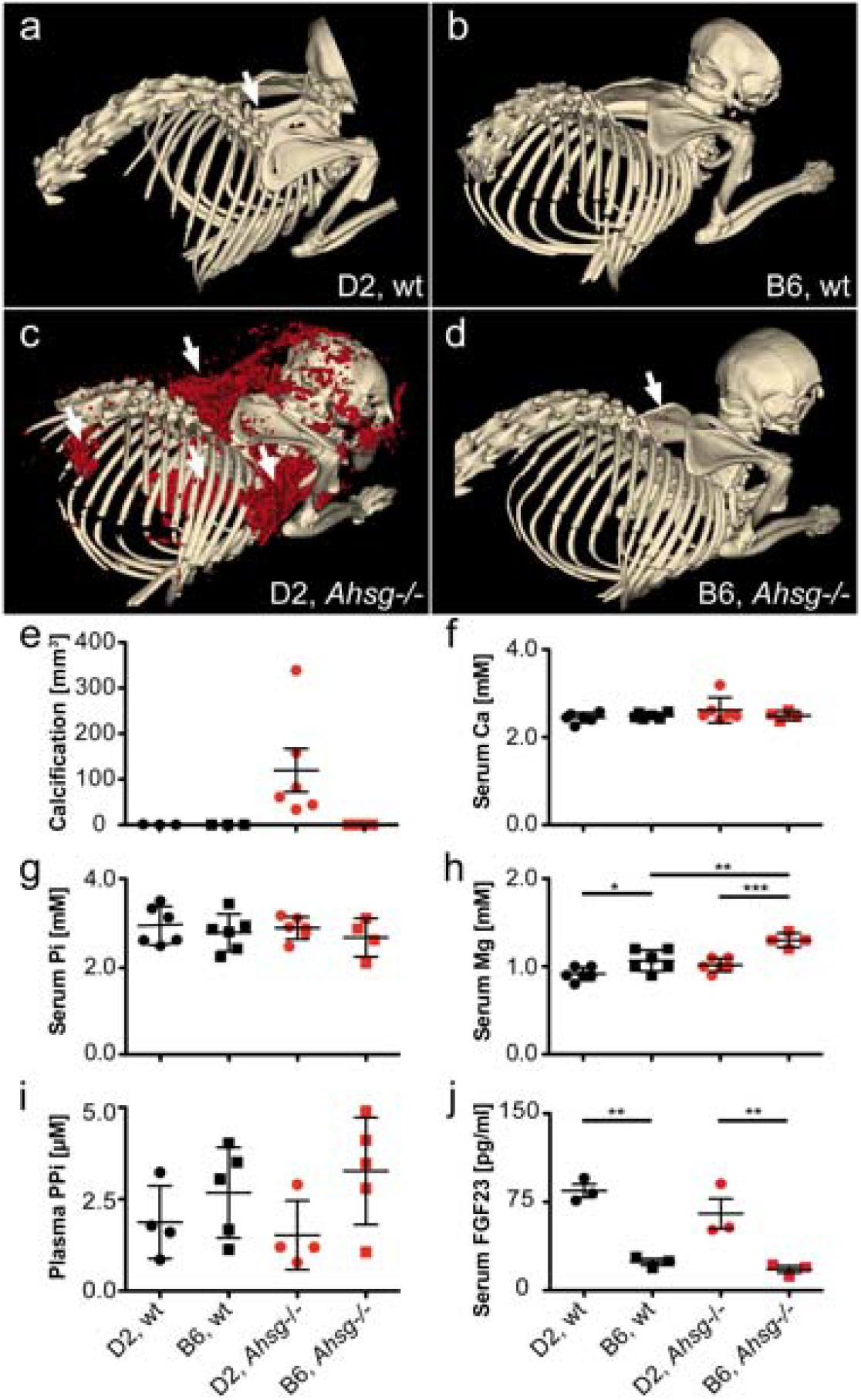
Extensive soft tissue calcification and hypomagnesemia at age 3 months in fetuin-A deficient mice maintained against the genetic background DBA/2. a, b, Wildtype (wt) and c, d, fetuin-A-deficient (*Ahsg−/−*) mice maintained against the genetic background DBA/2 (D2) or C57BL/6 (B6) were analyzed by computed tomography, and calcified lesions were segmented (red color) and e, quantified. Arrows point to small calcified lesions in a, the interscapular brown fat tissue of D2,wt and d, B6,*Ahsg−/−* mice. c, Arrows depict from left to right, massive calcified lesions in the left kidney, the heart, the interscapular brown adipose tissue BAT, and the axillary skin. f-i, Serum chemistry of calcification related electrolytes demonstrated f, normal serum total calcium, g, normal serum phosphate Pi, h, hypomagnesemia in D2,wt mice compared to B6,wt mice, and failure to induce serum magnesium in D2, *Ahsg−/−* compared to B6, *Ahsg−/−*, resulting in functional hypomagnesemia, i, plasma pyrophosphate, PPi was highly variable, yet slightly elevated in B6 mice compared to D2 mice and j, plasma FGF23 was significantly elevated in D2 compared to B6 mice, regardless of fetuin-A genotype. One-way ANOVA with Tukey’s multiple comparison test for statistical significance, *p<0.05, **p<0.01, ***p<0.001.

Recently, we showed that ectopic calcification in DBA/2 fetuin-A deficient mice starts in the microvasculature and is associated with premature ageing and cardiac failure, and that cellular osteogenesis was not involved [21].Thus, we concentrated on major extracellular regulators of calcification. Figure 1f,g show that serum calcium (Ca) and phosphate (Pi), major ionic drivers of calcification, were similar in DBA/2 and C57BL/6 wt and fetuin-A deficient mice. In contrast, serum magnesium (Mg), a potent inhibitor of calcification, was generally lower in DBA/2 mice confirming previous publications [17, 23] (Fig. 1h). C57BL/6 fetuin-A deficient mice, which do not spontaneously calcify, had increased serum magnesium levels compared to their wildtype littermates. In contrast fetuin-A deficient DBA/2 mice, which severely calcify, did not display elevated serum magnesium levels compared with wildtype littermates. Thus, an adaptive induction of serum Mg appears compromised in DBA/2 mice. Plasma levels of inorganic pyrophosphate (PPi), another systemic calcification inhibitor, were slightly elevated in C57BL/6 mice compared to DBA/2 mice independent of fetuin-A genotype (Fig. 1i). Serum FGF23 levels were significantly elevated in DBA/2 mice compared to C57BL/6 mice regardless of their fetuin-A genotype suggesting that the mice suffered from mineral dysbalance despite apparently normal serum phosphate (Fig. 1j). Taken together our results suggest that DBA/2 fetuin-A knockout mice have combined deficiencies of three potent circulating inhibitors of calcification, namely fetuin-A, Mg and PPi.

### Parenteral supplementation of fetuin-A reduced ectopic calcification development

We next asked if supplementation of the circulating inhibitors would affect the strong calcification phenotype of fetuin-A deficient DBA/2 mice. First, we injected intraperitoneally (i.p.) 0.24 g/kg bodyweight purified bovine fetuin-A five times weekly for a total of three weeks and then analysed soft tissues by histology, tissue chemistry, and X-ray analysis. Figures 2a-i show that saline injected fetuin-A deficient DBA/2 mice had severe dystrophic calcification in brown adipose tissue with lesions ranging from about 20 µm to several hundred µm in diameter (Fig. 2a), few small roundish calcified lesions in kidney and lung (Fig. 2b,c) and extensive fibrosing calcified lesions in the heart (Fig. 2d). Fetuin-A injection i.p. greatly reduced the number and size of calcified lesions (Fig. 2e-h, Suppl. Fig. S2) as well as the tissue calcium (Fig. 2i).

**Figure 2.**
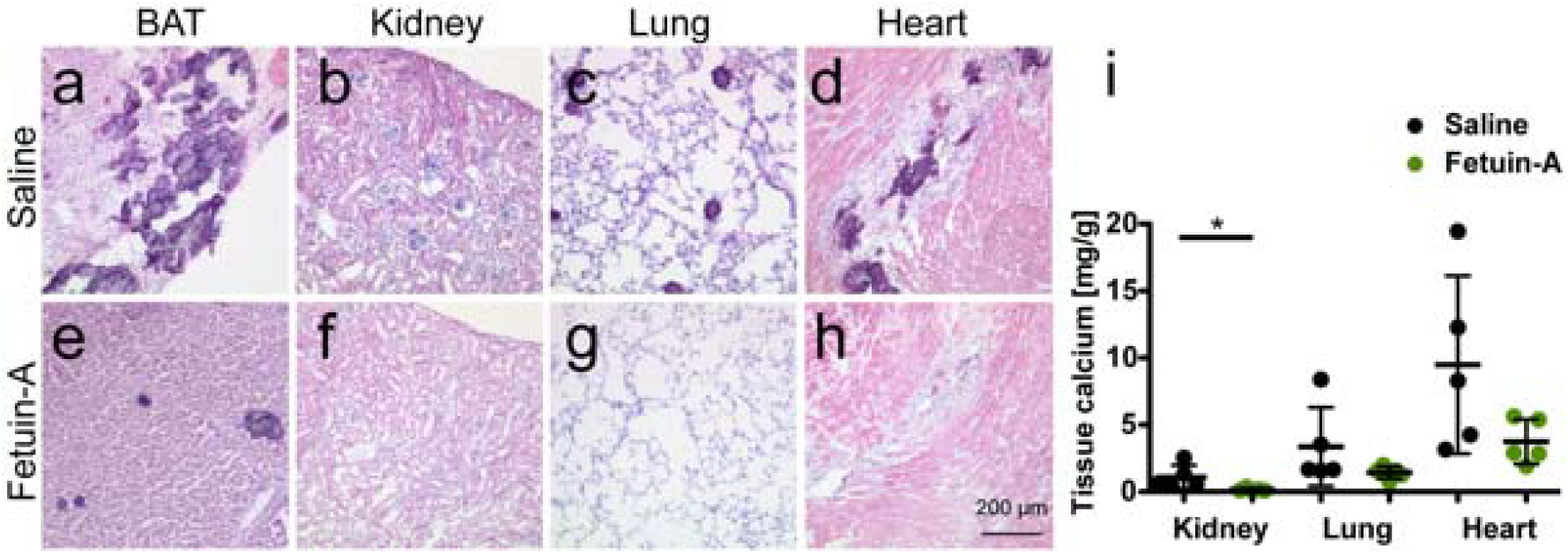
Fetuin-A supplementation attenuates soft tissue calcification in D2, *Ahsg−/−* mice. **a-d**, Three-week-old mice were injected injected intraperitoneal (i.p.) five times a week for three weeks, with saline or **e-h**, with 0.24 g/kg bodyweight fetuin-A. Shown are HE stained cryosections of **a,e** brown adipose tissue (BAT), **b,f**, kidney, **c,g**, lung, and **d,h** heart. Scale bar indicates 200 µm. Saline treated mice had numerous calcified lesions in their BAT, occasional lesions in kidney and lung, and extended fibrosing calcified lesions in myocard. In contrast, fetuin-A treated mice did not develop detectable calcified lesions in the same organs. **i**, Tissue calcium content was measured following chemical extraction of kidney, lung and heart of the animals. Treatment with fetuin-A strongly/markedly reduced the amount of tissue calcium in all analyzed organs. Student’s t-test was used to test for statistical significance, *p<0.05.

### Magnesium supplementation inhibits spontaneous soft tissue calcification

CT volume analysis of calcified lesions showed that dietary Mg supplementation strongly attenuated soft tissue calcification. DBA/2 mice on normal chow (0.2% Mg) developed 99 ± 36 mm^3^ calcifications within the first 11 weeks postnatally (Fig. 3a,b,e), while mice fed 1% Mg chow from postnatal week 3 onward developed 14 ± 36 mm^3^ calcifications (Fig. 3c,d,e). Chemical tissue calcium analysis and von Kossa histology confirmed that feeding 1 % Mg instead of 0.2 % Mg decreased calcifications in all organs (Fig. 3f-n). Analysis of major renal and gastrointestinal magnesium transporters showed that *Trpm7* expression was similar in kidney and gut tissue of all mice irrespective of genotype. In contrast, renal *Trpm6* expression was elevated in fetuin-A-deficient C57BL/6, but not in DBA/2 mice (Fig. 3o-r), suggesting that *Trpm6* gene induction mediates the elevated serum magnesium in fetuin-A deficient in C57BL/6, but not in DBA/2 mice (Fig. 1f).

**Figure 3.**
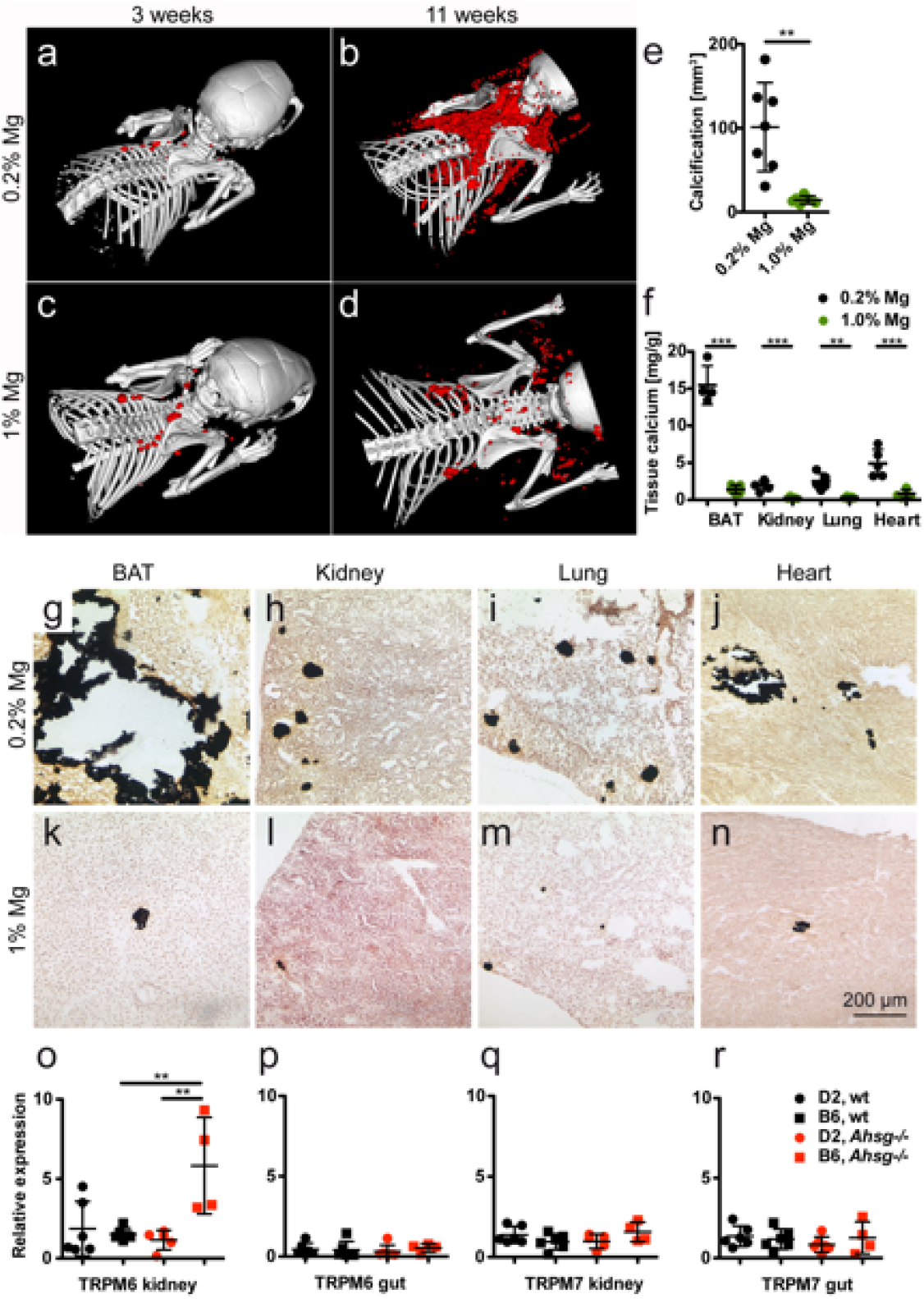
Dietary magnesium supplementation attenuates soft tissue calcification in D2, *Ahsg−/−* mice. **a-d**, Three-week-old mice were fed for eight weeks normal chow containing 0.2 % magnesium, or a high magnesium diet containing 1 % magnesium. Computed tomography of upper torso shows sparse calcified lesions in mice at the start of the feeding experiment (**a,c**), massive calcification after eight weeks on normal magnesium chow (**b**), and strongly attenuated calcification after eight weeks on high magnesium chow (**d**). **e**, Calcified tissue volumes determined by CT and segmentation. **F**, Tissue calcium content was measured following chemical extraction of brown adipose tissue (BAT), kidney, lung and heart. **g-n** show von Kossa stained cryosections of **g,k**, BAT, **h,l**, kidney, **I,m**, lung, and **j,n**, heart. Scale bar indicates 200 µm. Mice on normal chow had large calcified lesions in their BAT, occasional lesions in kidney and lung, and fibrosing calcified lesions in myocard. In contrast, mice fed with high magnesium developed sparse calcified lesions in the same organs. **o-r**, Unlike mice with the calcification resistant genetic background B6, mice with the calcification-prone genetic background D2 failed to induce *TRPM6* expression in the kidney, which resulted in the functional hypomagnesemia described in Fig. 1. Student t-test and one-way ANOVA with Tukey’s multiple comparison test for statistical significance, *p<0.05, **p<0.01, ***p<0.001.

### Dietary phosphate levels have a direct impact on calcification severity

Elevated serum FGF23 levels in DBA/2 mice compared to C57BL/6 suggested dysregulated phosphate handling despite apparently normal serum phosphate, a situation reminiscent of many CKD patients. We therefore studied the influence of dietary phosphate (high phosphate, 0.8% vs. low phosphate, 0.2%) on the calcification phenotype. CT analysis showed that mice on normal chow (0.4% Pi) developed 99 ± 36 mm^3^ calcifications within 11 weeks postnatal (Fig. 4a,b,g), while mice fed 0.2% Pi chow from week 3 postnatal onward developed 7 ± 12 mm^3^ calcifications (Fig. 4e,f,g). Feeding 0.8% Pi increased the volume of calcified lesions to 318 ± 130 mm^3^ (Fig. 4c,d,g). Tissue calcium analysis and von Kossa histology confirmed these findings in kidney, lung and heart, but not in brown adipose tissue. Feeding 0.8% Pi increased both the size of the lesions and the tissue calcium content in all organs analysed (Fig. 4h-t).

**Figure 4.**
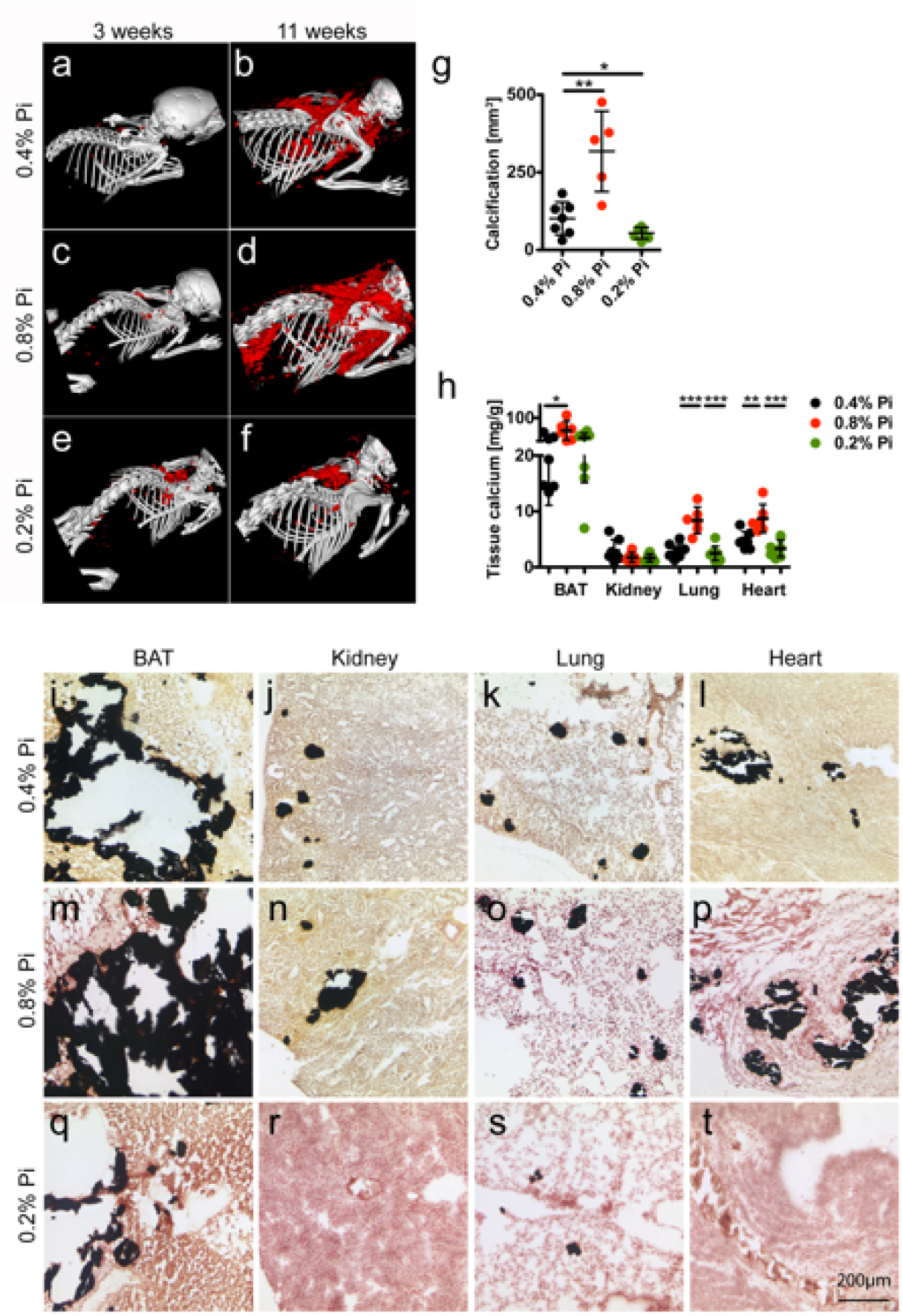
Dietary phosphate controls soft tissue calcification in D2, *Ahsg−/−* mice. **a-f**, Three-week-old mice were fed for eight weeks with normal chow containing 0.4% phosphorous (Pi) or with a low or high phosphate diet containing 0.2% and 0.8% phosphorous, respectively. Computed tomography of upper torso shows sparse calcified lesions in mice at the start of the feeding experiment (**a,c,e**). After feeding 8 % Pi the animals developed markedly more calcified lesions than control animals (**b,d**), while animals on low phosphate diet (0.2%) show strongly attenuated calcification (**f**). **g**, Calcified tissue volumes determined by CT and segmentation. **h**, Tissue calcium content was measured following chemical extraction of brown adipose tissue (BAT), kidney, lung and heart. **i-t** show von Kossa stained cryosections of **i,m,q**, BAT, **j,n,r**, kidney, **k,o,s**, lung, and **l,p,t**, heart. Scale bar indicates 200 µm. **i-l**, Mice on 0.4 % Pi chow had large calcified lesions in their BAT, occasional lesions in kidney and lung, and fibrosing calcified lesions in myocard. **m-p**, Mice on high 0.8% Pi diet had increased calcified lesions and **q-t**, mice on low 0.2% Pi diet had reduced calcified lesions in all organs studied. One-way ANOVA with Tukey’s multiple comparison test for statistical significance, *p<0.05, **p<0.01, ***p<0.001.

### Parenteral pyrophosphate inhibits the formation of ectopic calcification

Next we asked if supplementation of pyrophosphate would affect the calcification phenotype of DBA/2 mice. Starting at postnatal week 3, we injected daily for eight weeks, i.p. boluses of 0.10 g/kg bodyweight sodium pyrophosphate. CT showed that fetuin-A deficient DBA/2 mice injected with saline developed 91 ± 17 mm^3^ interscapular calcifications within the first 11 weeks postnatal (Fig. 5a,c), while age matched mice injected with pyrophosphate developed 9 ± 3 mm^3^ calcifications (Fig. 5b,c). Chemical tissue calcium analysis and von Kossa histology confirmed that pyrophosphate injection decreased calcifications in all organs analysed (Fig. 4d-l). Notably, pyrophosphate-injected fetuin-A deficient DBA/2 mice had rare calcified lesions in brown adipose tissue, a tissue most prone to calcification. The calcification phenotype of pyrophosphate treated mice was similar to untreated C57BL/6 mice (Fig. 1d), suggesting that pyrophosphate deficiency in DBA/2 mice is a major determinant of dystrophic calcification.

**Figure 5.**
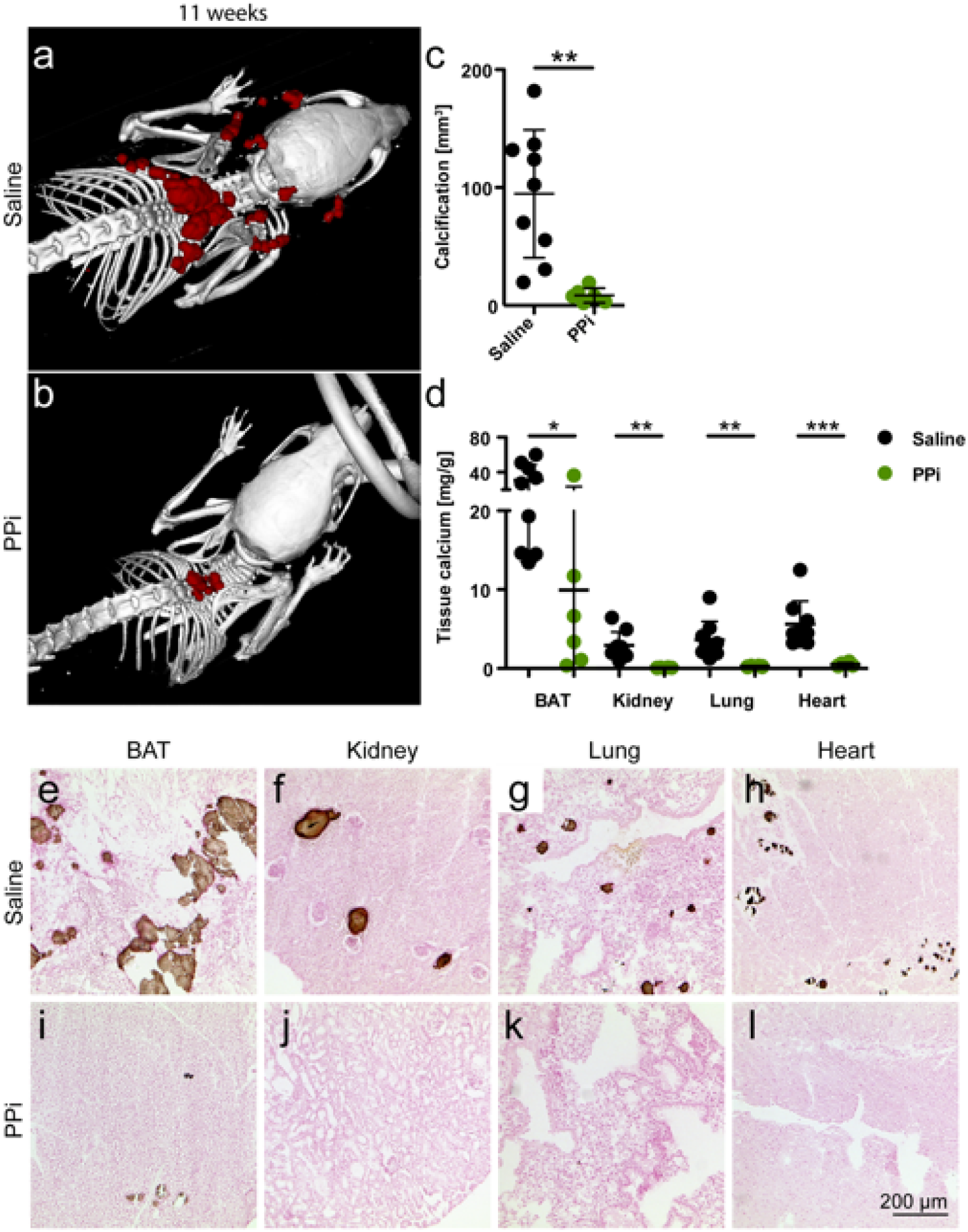
Pyrophosphate supplementation attenuates soft tissue calcification D2, *Ahsg−/−* mice. **a,d,e-h** Three-week-old mice were injected daily for eight weeks with saline or **b,d,i-l**, with 0.10 g/kg bodyweight sodium pyrophosphate (PPi). Computed tomography of upper torso shows massive calcification after eight weeks of saline injection (**a**), and strongly attenuated calcification after eight weeks of PPi injection (**b**). **c**, Calcified tissue volumes determined by CT and segmentation. **d**, Tissue calcium content was measured following chemical extraction of brown adipose tissue (BAT), kidney, lung and heart. **e-l** show von Kossa stained cryosections of **e,i**, BAT, **f,j**, kidney, **g,k**, lung, and **h,l**, heart. Scale bar indicates 200 µm. **a,e-h**, Saline treated mice had numerous calcified lesions in their BAT, occasional lesions in kidney and lung, and fibrosing calcified lesions in myocard. **b,i-l**, In contrast, PPi treated mice had reduced calcified lesions only in interscapular BAT. Student t-test for statistical significance, *p<0.05, **p<0.01, ***p<0.001.

### Fetuin-A and Abcc6 deficiency are the main calcification risk factors in D2, *Ahsg-/-* mice

Next we studied the expression of Abcc6, an ATP cassette transporter protein involved in extracellular pyrophosphate metabolism [24, 25]. The single nucleotide polymorphism (SNP) rs32756904 signifying a 5-bp deletion in the Abcc6 transcript [26], was readily detected by differential PCR and DNA gel electrophoresis in DBA/2 mice independent of the fetuin-A genotype (Fig. 6a). Supplementary figure S3 shows the G>A mutation flanking the splice donor site of exon 14 of the murine *Abcc6* gene of DBA/2 mice resulting in reduced mRNA expression levels in liver and kidney, the two major Abcc6 expression sites (Fig. 6c,d). Immunoblotting confirmed reduced Abcc6 protein expression in livers of DBA/2 mice compared to C57BL/6 mice, and furthermore, failure to induce *Abcc6* expression in fetuin-A deficient mouse strains. The SNP illustrated in Supplementary Figure S3 was detected in all DBA/2 mice, but never in C57BL/6 mice, suggesting that the hypomorphic G>A mutation added to the overall calcification risk in DBA/2 mice.

**Figure 6.**
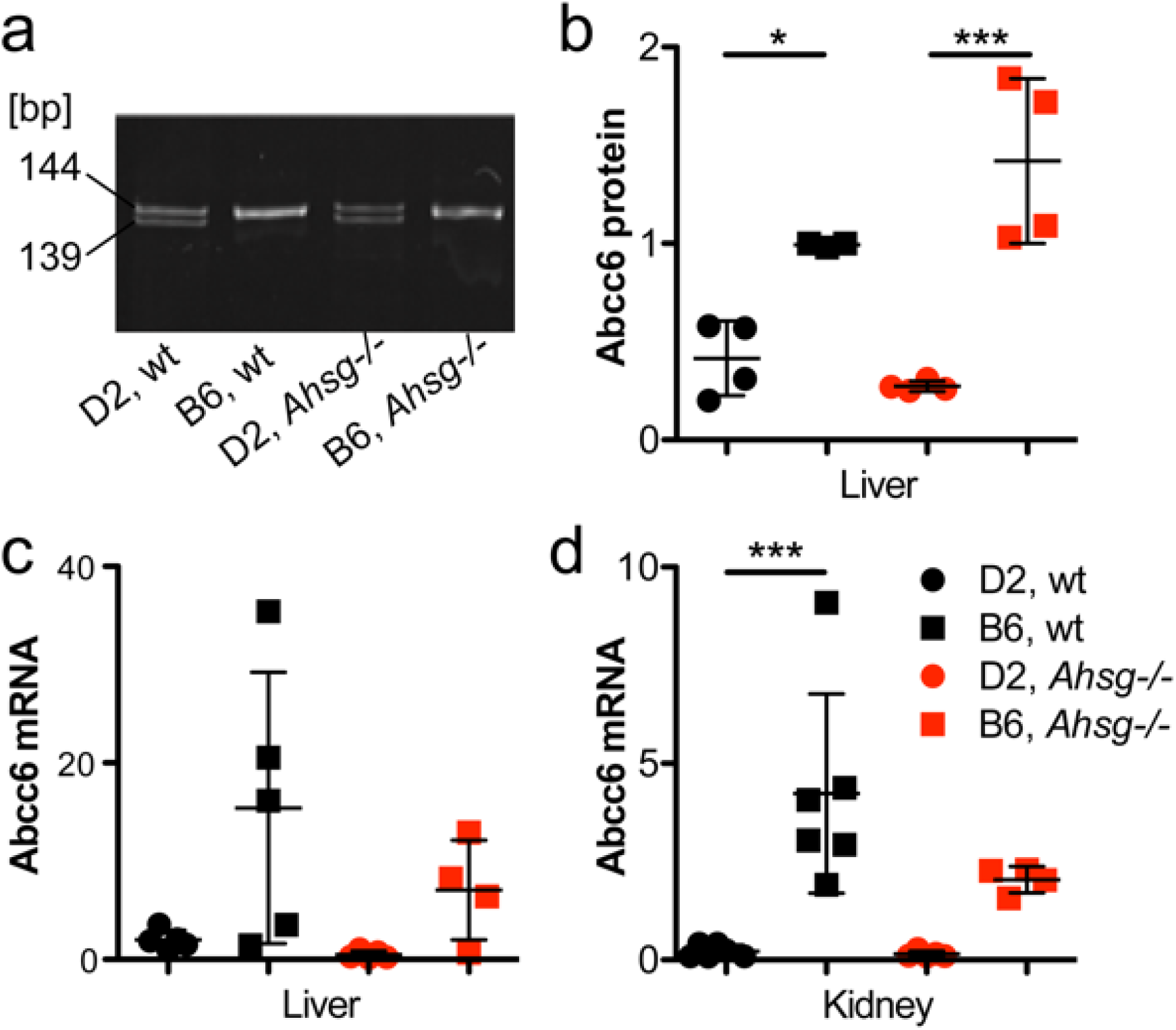
A single nucleotide polymorphism in the gene Abcc6, encoding the ATP-binding cassette subfamily C member 6, a regulator of serum PPi levels in DBA/2 mice is associated with hypomorphic Abcc6 expression in liver and kidney. a, SNP rs32756904 causing a 5-bp deletion in the Abcc6 transcript was detected by differential PCR and DNA gel electrophoresis. Amplicon size is given in base pairs [bp]. b, The polymorphism is associated with reduced expression of correctly spliced Abcc6 transcript, which was verified by immunoblotting of Abcc6 protein in the liver, c, d, and by qPCR of mRNA in liver and kidney, respectively. Overall the hypomorphic Abcc6 mutation results in reduced steady state extracellular pyrophosphate, a potent inhibitor of calcification. For a genomic analysis of the hypomorphic Abcc6 mutation see supplemental figure S3 online. One-way ANOVA with Tukey’s multiple comparison test for statistical significance, *p<0.05, **p<0.01, ***p<0.001.

To confirm a major role of *Abcc6* and thus of pyrophosphate metabolism in the strong calcification phenotype of fetuin-A deficient DBA/2 mice, we analysed by linkage analysis the contribution of this hypomorphic Abcc6 mutation to the overall calcification risk. To this end we crossed fetuin-A deficient DBA/2 male with C57BL/6 female mice. The heterozygous F_1_ offspring were randomly intercrossed to create an F_2_ generation (Fig. 7a). We analyzed calcification in 177 F_2_ offspring (86 males, 91 females) by CT and stratified the mice with respect to calcification: **I**, no detectable calcified lesions, 65 animals; **II**, single calcified lesion in brown adipose tissue, 42 animals; **III**, two clearly separated lesions in brown adipose tissue, 42 animals; **IV**, calcified lesions in brown adipose tissue, myocardium and adrenal fat pad, 24 animals; **V**, calcified lesions throughout the body, 4 animals (Fig. 7b). Heterozygous F_1_ progeny had at most one calcified lesion in brown adipose tissue, indicating that half the gene dosage of calcification regulatory genes from resistant C57BL/6 genetic background was sufficient to prevent widespread calcification even in the absence of fetuin-A. Figure 7b illustrates that F_2_ mice carrying homozygous the mutant A allele of *Abcc6* had severe calcifications (III, IV, V; with one exception in category II). Mice with the homozygous wildtype G allele had no detectable calcified lesions. Mice carrying both the mutant and the wildtype allele had variable calcification including categories I, II, III, IV. This result suggested a necessary, but not an exclusive determination by the SNP rs32756904 of the strong calcification phenotype of fetuin-A deficient mice. Presently we cannot offer a similar frequency plot regarding *Trpm6* genetics, but the clear segregation of the mice with little or no calcification (I, II) and the mice with severe calcification (III, IV, V) with no overlap suggested that few if any additional genes beyond *Fetua* / *Ahsg, Abcc6*, and *Trpm6* determined the calcification propensity.

**Figure 7.**
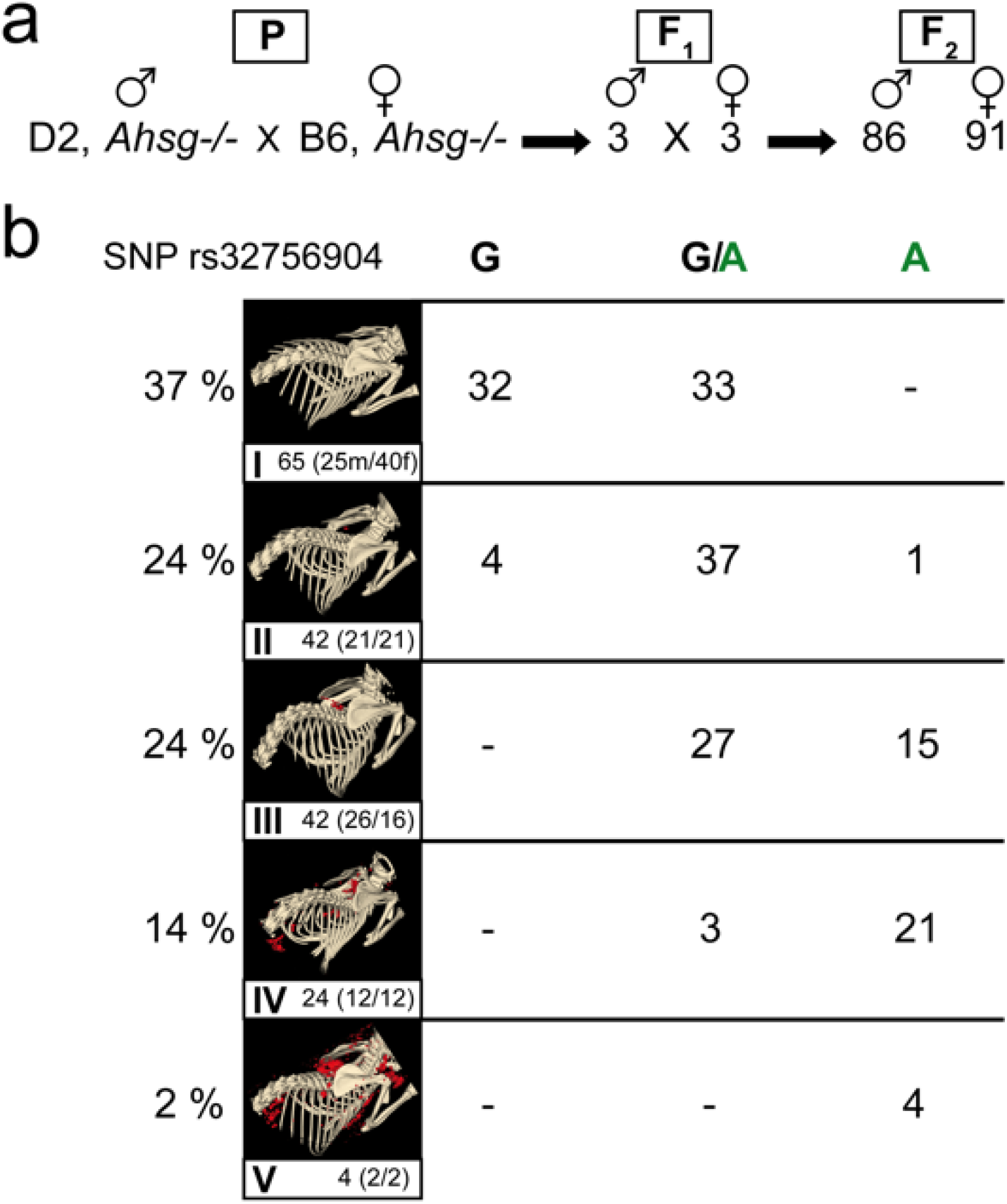
Linkage analysis of SNP rs32756904 with calcification phenotype in progeny of fetuin-A deficient DBA/2 and C57BL/6 mice. a, One male D2, *Ahsg−/−* mouse was naturally mated with one female B6, *Ahsg−/−* mouse (parental generation – P). b, Three breeding pairs of their hybrid progeny (F_1_) produced 177 F_2_-offspring (86 male, 91 female), which were genotyped by PCR and genomic sequencing for the hypomorphic Abcc6 single nucleotide polymorphism rs32756904. Calcification was individually scored by computed tomography and segmentation of non-skeletal mineralized tissue (red color). Calcification was scored in five categories ranging from no calcified lesions detected (I), one calcified lesion detected in the brown adipose tissue BAT (II), more than one calcified lesion detected in BAT, but not in other tissues (III), numerous calcified lesions detected in BAT, kidney and myocard (IV), and numerous massive calcified lesions in BAT, kidney, myocard and axillary skin (V). The clear segregation of scores with SNP rs32756904 suggests that a limited number of 1-2 additional genes besides fetuin-A deficiency and SNP rs32756904 determine the severity of the calcification phenotype of the mice.

## Discussion

To prevent crystallization and deposition of calcium mineral in tissues not meant to mineralize, mineralization regulators evolved. Fetuin-A is a potent systemic inhibitor of ectopic calcification mediating the solubilization and clearance of circulating protein-mineral complexes, which might otherwise cause calcification [27, 28]. In this present study parenteral supplementation of fetuin-A inhibited the formation of ectopic calcifications in our mouse model. In humans, complete fetuin-A deficiency was not reported until recently despite the fact that human fetuin-A/a2HS glycoprotein was a commonly used paternity marker [29]. Partial fetuin-A deficiency was, however, reported in many studies of CKD, and correlated with increased serum calcification propensity [30–32]. Recently, genetic fetuin-A deficiency was found associated with infantile cortical hyperostosis (Caffey disease) [33], a pathology compatible with the post-weaning epiphysiolysis causing distal femur dysplasia and foreshortened hindlimbs in fetuin-A-deficient mice maintained against the genetic background C57BL/6 [34]. Unlike the fetuin-A deficient mice maintained against the background DBA/A studied here, these mice had no overt calcification outside the skeleton, suggesting that growth plate cartilage, a tissue physiologically meant to calcify during bone development, readily calcifies, when fetuin-A is lacking, while other tissues require additional calcification risk factors. Indeed, this is the main finding of this current work.

We showed that DBA/2 mice had hypomagnesemia regardless of their fetuin-A genotype due to altered expression of magnesium transporter *Trpm6.* The protective action of Mg in calcification is known [35], as is its basic regulation [36]. The adaptive regulation of Mg in calcification however, is only partially understood. We hypothesized that DBA/2 mice are more prone to calcification than are C57BL/6 mice, because of renal Mg wasting [37] and thus loss of systemic Mg, a potent crystal nucleation inhibitor [38]. A recent study showed however, that kidney-specific TRPM6 knockout mice had no phenotype in contrast to intestine-specific knockout [39]. These results recapitulated clinical studies with patients with TRPM6 null mutations clearly showing that a defect in intestinal Mg2+ uptake causes hypomagnesemia. In any case, hypomagnesemia compounds the deficiency of serum fetuin-A, a crystal growth inhibitor. Magnesium supplementation prevented calcification in a mouse model of pseudoxanthoma elasticum (PXE), *Abcc6*−/− mice [14], and reduced nephrocalcinosis [40] as well as vascular and soft tissue calcifications in a CKD rat model [41]. Serum Mg is considered a poor marker of Mg deficiency and is not routinely monitored in CKD patients [35]. However, in CKD patients Mg containing phosphate binders, supplementation with Mg-oxide or increased Mg in dialysate reduced serum calcification propensity [42] and/or the progression of cardiovascular calcifications [43]. These findings are in line with our observation that magnesium supplementation prevented ectopic calcification in DBA/2 mice. Hypomagnesemia in humans is common in about 15% of the general population. Inherited hypomagnesemia is rare [44] but malnutrition and acquired metabolic disease seem to account for most cases. Drugs such as proton pump inhibitors also reduce serum magnesium [45, 46]. When hypomagnesemia combines with calcification risk factors including CKD, it might accelerate the development of ectopic calcification. Nevertheless in CKD patients, Mg supplementation must be tightly monitored to avoid hypermagnesemia, despite reports of it being safe [42].

DBA/2 mice regardless of their fetuin-A genotype had hypomagnesemia associated with elevated serum FGF23. Likewise, rats on magnesium-deficient diet had elevated serum FGF23 [47] and CKD cats had inverse association of serum Mg and FGF23 independent of serum phosphate [48]. Collectively these results suggest that serum FGF23 may be associated with hypomagnesemia in addition to the well-established association with hyperphosphatemia and morbidity. CKD-associated hyperphosphatemia is a result of impaired renal excretion and is directly associated with cardiovascular calcification [49, 50]. Additionally, elevated phosphate levels might promote coronary atheroma burden in non-uremic patients [51]. Thus, dietary phosphate restriction is paramount in dialysis patient care [52]. Although several studies report on the effectiveness of phosphate binders on calcification inhibition in different animal studies [53–55], the effect of dietary phosphate restriction on calcification has been studied in surprisingly little detail [56, 57]. The evaluation of dietary phosphate restriction in CKD patients remains challenging as dietary phosphate restriction generally implies protein restriction and thus potential malnutrition.

Dystrophic calcification in genetically predisposed mouse strains such as C3H/He, DBA/2, BALB/c and 129S1 is a complex genetic trait still not fully understood. Genetic mapping identified a single major locus, designated *Dyscalc1*, located on proximal chromosome 7 (chr 7), and three additional modifier loci associated with dystrophic cardiac calcinosis (DCC) following myocardial injury [58]. Integrative genomics identified the gene mapping to the *Dyscalc1* locus as *Abcc6*, the ATP-binding cassette C6 [26]. Abcc6 is a membrane transporter with hitherto unknown substrate specificity. Forced expression of Abcc6 was associated with increased cellular release of ATP, which rapidly breaks down into AMP and the potent calcification inhibitor pyrophosphate PPi [24, 25, 59]. PPi is a calcium phosphate crystal nucleation and growth inhibitor that interferes with crystal lattice formation. Circulating PPi is mostly liver-derived and relatively long-lived [25]. In hemodialysis patients, low PPi levels [60] are associated with increased calcification risk [61]. We found reduced plasma PPi levels in D2 mice compared to B6 mice, which may result from reduced Abcc6 expression in D2 mice. Recent studies showed that plasma PPi deficiency is the major, if not the exclusive cause of dystrophic calcification in *Abcc6−/−* mice [62]. A non-coding splice variant of Abcc6 was deemed responsible for the PPi deficiency in calcification prone C3H/He mice. A single nucleotide polymorphism rs32756904 encodes an additional splice donor site and thus the hypomorphic splice variant of Abcc6. We detected rs32756904 in all of our DBA/2 mice, underscoring the importance of genetic background in the expression of dystrophic calcification [63, 64].

Hypomorphic mutations in the *Abcc6* gene are deemed responsible for both dystrophic calcification in mice and for PXE in humans [26]. Children suffering from generalized arterial calcification of infancy (GACI), caused by a mutation of the gene *Enpp1*, which is responsible for the conversion of ATP into AMP and PPi, could also benefit from PPi supplementation therapy. In addition to PXE and GACI, Hutchinson-Gilford progeria syndrome (HGPS), was linked to reduced PPi levels associated with the development of extracellular matrix calcification [13]. Although mouse models for HGPS, GACI and PXE show promising results when treated with PPi, human studies remain to be conducted [65].

In conclusion, we describe that a compound deficiency of fetuin-A, magnesium and pyrophosphate in DBA/2 fetuin-A knockout mice leads to arguably one of the most severe soft tissue calcification phenotypes known. We showed that dietary magnesium, low dietary phosphate, parenteral fetuin-A and pyrophosphate supplementation all prevented the formation of calcified lesions in fetuin-A deficient DBA/2 mice suggesting that similar therapeutic approaches might also reduce early stage CKD-associated cardiovascular calcifications and in calciphylaxis. Notably, these calcifications occurred without any signs of osteogenic conversion of calcifying cells, stressing the importance of extracellular mineral balance to prevent unwanted mineralization [21].

## Materials and Methods

### Animals

All animal experiments were conducted in agreement with German animal protection law and were approved by the state animal welfare committee. Wildtype (wt) and fetuin-A deficient (*Ahsg−/−*) mice on DBA/2N (D2) and C57BL/6N (B6) genetic background (ILAR entries D2-*Ahsg*^tm1wja^ and B6-*Ahsg*^tm1wja^ according to ILAR nomenclature) were maintained in a temperature-controlled room on a 12-hour light/dark cycle. Chow (ssniff^®^ R/MH, V1535-0, Soest, Germany) and water were given ad libitum. Mice were kept at the animal facility of RWTH Aachen University Clinics.

For gene segregation and linkage analysis, one D2, *Ahsg−/−* male was mated with one B6, *Ahsg−/−* female. F1 offspring was intercrossed, and 177 F2 offspring were obtained.

### Supplementation therapy

DBA/2 mice deficient for fetuin-A were enclosed in the experiment at three weeks of age. The study included five treatment groups (n≥6 per group) receiving either 0.24 g/kg bodyweight bovine fetuin-A (Sigma) or 0.10 g/kg bodyweight PPi (Sigma) via daily intraperitoneal injections for 3 or 8 weeks, respectively or 1.0% magnesium, 0.2% Pi or 0.8% Pi via dietary supplementation for 8 weeks. Control animals received 0.9% NaCl via intraperitoneal injection and normal chow.

Mice were sacrificed with an overdose of isoflurane, exsanguinated, perfused with ice-cold PBS and organs were harvested. A piece of each of the following tissues (kidney, heart, lung, brown adipose tissue of the neck) was frozen in liquid nitrogen for calcium measurement, another piece was embedded in Tissue-Tek O.C.T. compound and snap frozen for histology.

### High-Resolution Computed Tomography

Mice were anaesthetized with isoflurane and placed in a high-resolution computed tomograph (Tomoscope DUO, CT-Imaging, Erlangen, Germany). The settings of the CT scan were: 65 kV / 0.5 mA, and 720 projections were acquired over 90 seconds. Analysis of the CT scans was performed with the Imalytics Preclinical Software [66]. Full-body acquisitions were obtained to examine in three dimensions the overall state of mineralization in the mice.

### Tissues

At the indicated ages mice were sacrificed with an overdose of isoflurane and exsanguinated. Animals were perfused with 20 ml ice-cold PBS to rinse blood from the circulation unless stated otherwise.

Calcium content in the tissue (kidney, lung, heart, brown adipose tissue) was determined using a colorimetric o-cresolphthalein based assay (Randox Laboratories, Crumlin, UK). Tissue samples were incubated overnight in 0.6 M HCl followed by homogenization with a Qiagen mixer mill. Homogenates were centrifuged for 10 minutes at 14000 rpm and supernatant was neutralized with ammonium chloride buffer. The subsequent assay was performed according to the manufacturer’s protocol.

### Clinical chemistry (Ca, Mg, P_i_, PP_i_) and FGF23 serum levels

Blood was clotted and centrifuged at 2000xg for 10 min. Serum was harvested and snap-frozen in liquid nitrogen. Standard serum chemistry (Ca, Mg, Pi) was commissioned by the Institute of Laboratory Animal Science at the University Hospital Aachen. Plasma PP_i_ was measured in microfiltered plasma as described [25]. Serum FGF-23 was measured with ELISA (Mouse Fibroblast growth factor 23 ELISA Kit, Cusabio, CSB-EL008629MO) according to manufacturers instructions, with samples diluted 1:2 in sample diluent.

### DNA-Analysis for SNP rs32756904 in the gene *Abcc6*

Genomic DNA was isolated from tail tip following established protocols. Using primers flanking the G > A mutation side (forward, TGGCCCACTCTTGTATCTCC /reverse, TTGGGTACCAAGTGACACGA) a 192 kDa PCR fragment was amplified and separated on a 1,5 % agarose gel. Amplicons were stained by ethidium bromide, excercised with an scalpel under UV light and afterwards purified by QIAquick^®^ Gel Extraction Kit (Qiagen). Sequencing of the PCR products was carried out by eurofins genomics.

### cDNA synthesis and quantitative real time PCR

Total RNA from the respective organ was isolated with GeneJet RNA Purification Kit (Thermo Fisher Scientific). Isolated RNA was treated with DNase I (Roche Diagnostics) to eliminate genomic DNA. Afterwards total RNA was reverse transcribed (Maxima First Strand cDNA Synthesis Kit, Thermo Fisher Scientific) into cDNA, which served as template both for sequencing (Eurofins Genomics) and for quantitative real-time PCR of three target genes (*Abcc6, TRMP6, TRPM7*). Supplementary Table S1 (online) shows the corresponding primer sequences. Primers were previously tested for efficiency. Fold changes were determined using the ΔΔCT method and glyceraldehyde 3-phosphate dehydrogenase *Gapdh* as the reference gene. Maxima SYBR Green/ROX qPCR Master Mix (Thermo Scientific) was used for qPCR and 20 ng of cDNA were added per reaction. Reaction specificity was determined by the dissociation curve.

By using primers (forward, CGAGTGTCCTTTGACCGGCT /reverse, TGGGCTCTCCTGGGACCAA) flanking the 5-bp deletion in Abcc6 cDNA of fetuin-A deficient mice on DBA/2N a 144-bp or 139-bp PCR fragment was amplified representing wildtype and mutant alleles, respectively. For the separation of the 5-bp shortened mutant fragment from the wildtype amplicon, high-resolution electrophoresis was performed using a 10% polyacrylamide gel with an acryl-amide/bisacryl-amide-relation of 19:1. Sequencing of PCR products was carried out by eurofins genomics.

### Immunoblotting

For Abcc6 Western blot analysis liver samples frozen in liquid nitrogen were powdered with a mortar. Using Lysis Buffer (10 mL/ g organ;150 mM NaCl, 20 mM Tris base pH 7.5, 1% Triton X-100, 1 mM EDTA, 1xRoche inhibitor cocktail (Roche Diagnostics GmbH)) the proteins were extracted for 30 minutes at 4°C by shaking. Cell fragments were separated by centrifugation (4°C and 20000xg for 15 min). The protein concentration of the supernatant was determined using a BCA protein assay reagent kit (Pierce). Protein samples (100 µg/lane) were separated using a gradient gel (NuPAGE® Novex 4-12% Bis-Tris Gel 1.0 mm, Thermo Fisher Scientific) and transferred onto nitrocellulose membrane (Amersham Protran, Sigma Aldrich) by wet electroblotting (Mini Trans-Blot^®^ Cell, Biorad) at 120 V for 2,5 h. The membrane was blocked with 3% nonfat dried milk powder in PBS-T (PBS supplemented with 0.05% Tween-20, Applichem) for 45 min at room temperature. Primary antibody (MRP6 (M6II-68) monoclonal rat anti-mouse Abcc6 [36] was diluted 1:1000 in 1% blocking solution and applied over night at 4°C. Secondary antibody (horseradish peroxidase coupled rabbit anti-rat IgG, Dako) was diluted 1:2500 in 1% blocking solution and incubated for 1 h at room temperature. Following antibody incubations membranes were washed three times with PBS-T for 5 min. Bound antibody was detected by chemiluminescence in substrate solution (0.1M TRIS/HCl, pH 8.5, 1.25mM 3-aminopthalhydrazide, 0.45mM p-coumaric acid, 0.015% hydrogen peroxide) using a fluorescence imager (Fuji LAS Mini 4000). Abcc6 protein levels were relatively quantified using Image J software (Rasband, http://imagej.nih.gov/ij/).

For serum fetuin-A analysis murine serum was separated by sodium dodecyl sulphate polyacrylamide gel elecrophoresis using mini gels (10% acrylamide, 5 × 8 × 0.1 cm^3^, BioRad, Hercules, USA). Protein transfer onto nitrocellulose membrane was by semi-dry electroblotting (Owl HEP-1, Thermo Scientific, Waltham, MA, USA) at 6 V for 60 min. The membrane was blocked with 5% nonfat dried milk powder in PBS-T for 30 min at 37°C. Primary antibody (AS386 polyclonal rabbit anti-mouse fetuin-A) was diluted 1:2500 in blocking solution and applied for 1 h at 37°C. Secondary antibody (horseradish peroxidase coupled swine anti-rabbit IgG, Dako) was diluted 1:5000 in blocking solution and incubated for 1 h at 37°C. Following antibody incubations membranes were washed three times with PBS-T for 5 min and bound antibody was detected by chemiluminescence as described above.

### Histology

6 µm thick sections were cut and fixed with zinc fixative (BD Biosciences, Heidelberg, Germany). Subsequently, H/E staining or von Kossa staining was performed. The sections were analyzed using a Leica DMRX microscope (Leica Microsystems GmbH, Wetzlar, Germany) and DISKUS software (Carl H. Hilgers, Technisches Büro, Königswinter, Germany).

### Statistics

All statistical analyses were carried out using GraphPad Prism version 5.0c and are given as mean values ± standard deviations. T-test was used for comparison of single treatments versus control group. One-way ANOVA with Tukey’s multiple comparison test was used to test for differences in non-sized matched experimental groups.

## Acknowledgements

This work was supported by grants awarded to WJD by the IZKF Aachen of the Medical Faculty of RWTH Aachen and by the German Research Foundation (DFG SFB/TRR219-Project C-03). We thank Koen van de Wetering (Jefferson Univ, Philadelphia, USA) for help with the pyrophosphate assay.

## Author Contributions Statement

ABa – performed experiments, analyzed data, drafted and revised the manuscript

CS – performed experiments, analyzed data, drafted and revised the manuscript

ABü – performed experiments, analyzed data, drafted and revised the manuscript

MH – designed study, revised the manuscript

FG – designed study, revised the manuscript

TG – provided material, revised the manuscript

WJD – designed study, analyzed data, drafted and revised the manuscript

## Conflict of interest statement

Authors declare no conflict of interest.

---

**Supplementary Figure S1.**
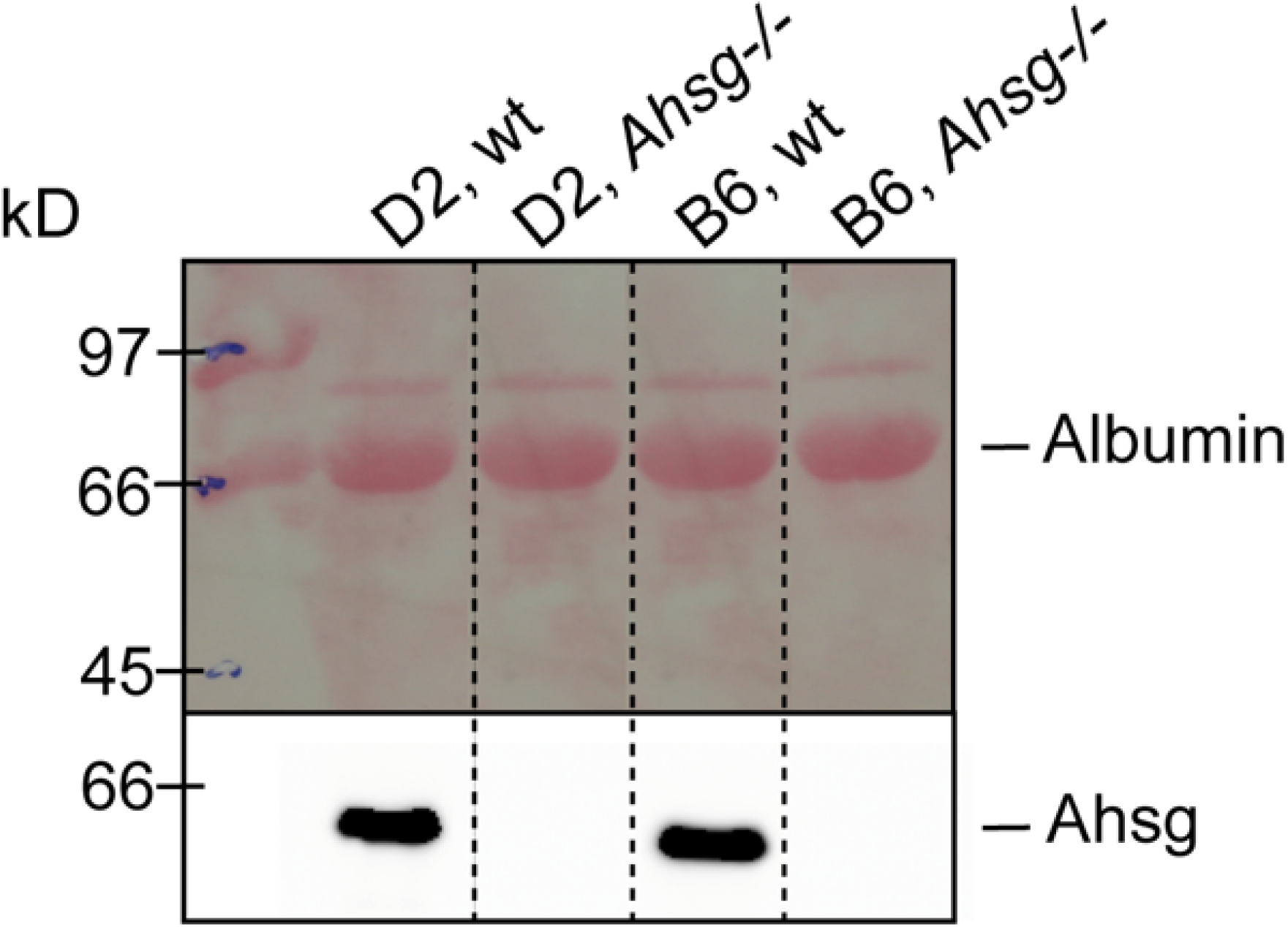
Serum immunoblot confirms complete lack of fetuin-A expression (*Ahsg*) in D2, *Ahsg−/−* and B6, *Ahsg−/−* mice.

**Supplementary Figure S2.**
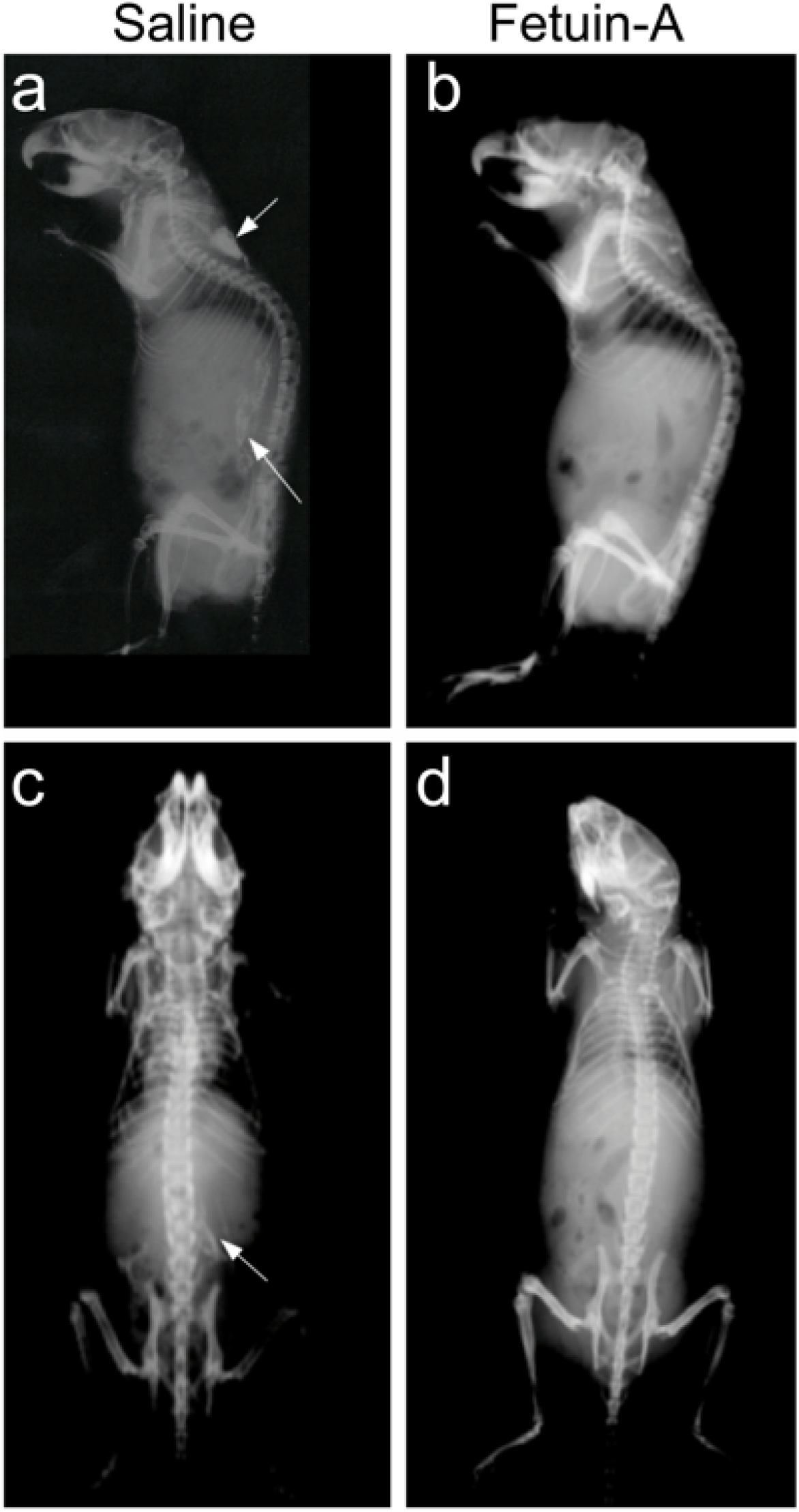
Fetuin-A supplementation attenuates soft tissue calcification in D2, *Ahsg*−/− mice. **a-d**, Three-week-old mice were injected i.p. five times a week for three weeks, with saline (**a,c**) or with 0.24 g/kg bodyweight fetuin-A (**b,d**). Lateral and dorsal radiographs show brown adipose tissue and renal fat calcification (arrows) in saline treated but not in fetuin-A treated mice.

**Supplementary Figure S3.**
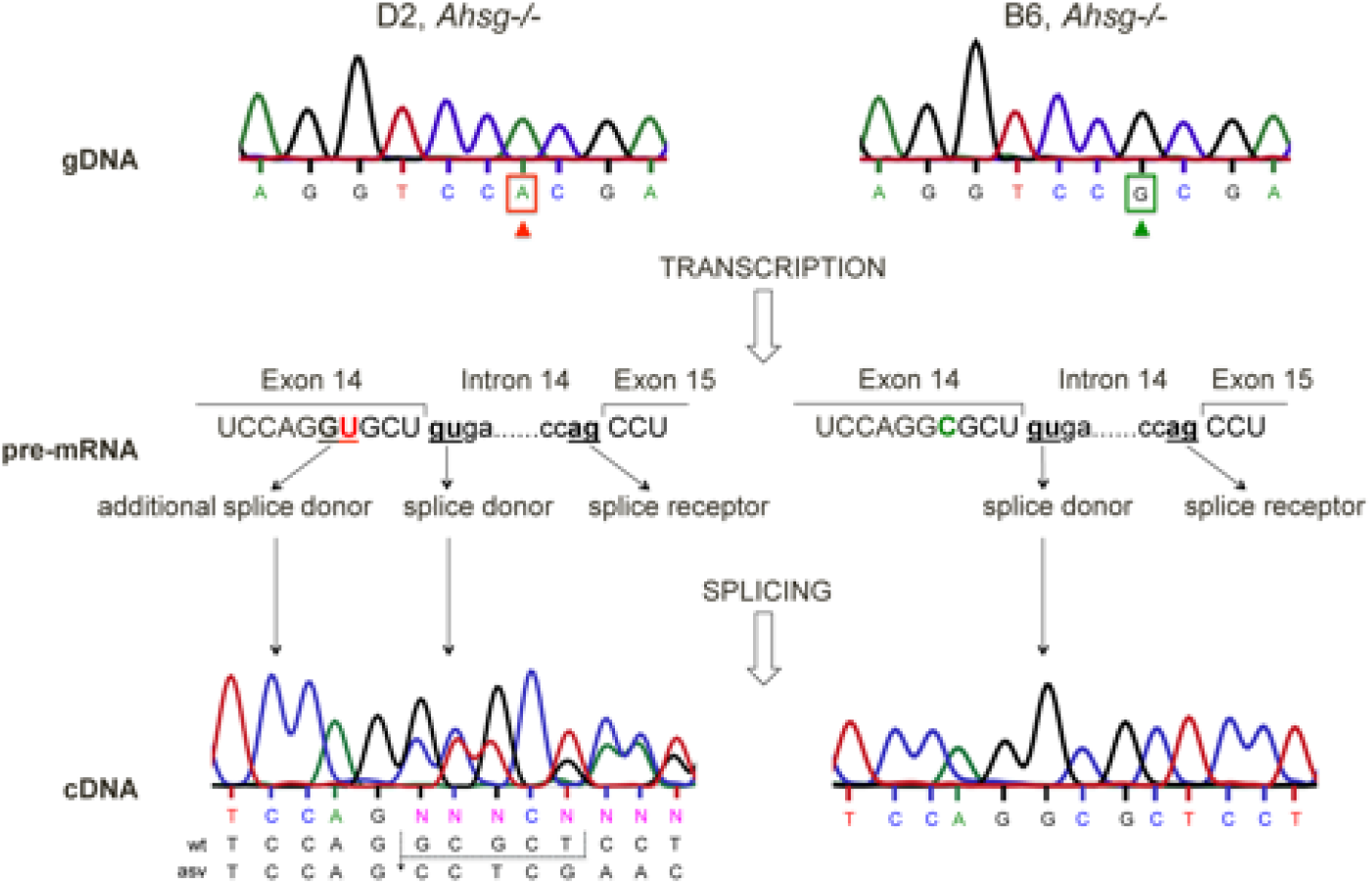
Genomic organization of SNP rs32756904 in the gene *Abcc6* on chromosome 7 in D2 and B6 fetuin-A deficient. B6 mice have unambiguous splice donor and acceptor sites in their genomic DNA (gDNA) resulting in a single pre-mRNA and a single mRNA transcript. In contrast, D2 mice carry a G>A mutation in their gDNA, creating an additional splice donor site five base pairs upstream of the wildtype splice acceptor site. This creates an alternative five base pairs shortened splice variant of the mRNA transcript, which is not translated. Thus, the hypomorphic SNP rs32756904 results in reduced expression of functional Abcc6 protein and reduced extracellular pyrophosphate levels.

**Supplementary Table S1.**
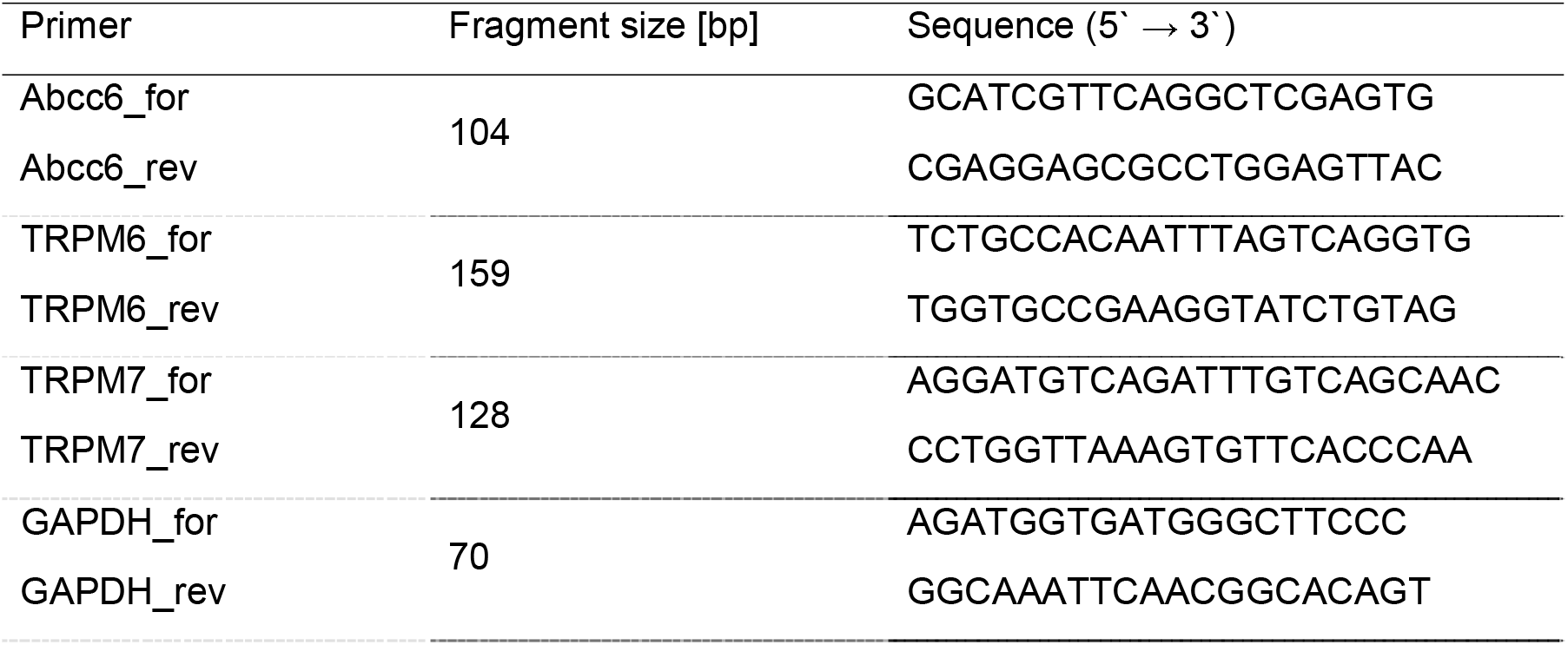
Primer sequences used for quantitative real-time PCR. Primers were designed using NCBI primer-BLAST or taken from Primer Bank. We ensured that all primers were spanning exon-exon junctions and primed the coding region of the gene.

## Notes

#### Summary of Updates

Title changed. Text changed in Abstract Results and Discussion for improved reading. Typos corrected. refs 33 , 34, 39, 43 added

